# The antiphage defense system CBASS controls resistance and enables killing by antifolate antibiotics in *Vibrio cholerae*

**DOI:** 10.1101/2023.02.27.530311

**Authors:** Susanne Brenzinger, Martina Airoldi, Adewale Joseph Ogunleye, Ana Rita Brochado

**Affiliations:** Department of Microbiology, Biocenter, University of Würzburg, Würzburg, Germany

## Abstract

Toxic bacterial modules, in particular toxin-antitoxin (TA) systems, have been long sought-after for their antimicrobial potential, although with limited success^1–6^. Here we show that the cyclic-oligonucleotide-based antiphage signaling system (CBASS), another example of a toxic module, increases sensitivity to well-established antifolate antibiotics, interferes with their synergy, and ultimately enables bacterial lysis by antifolates - classic bacteriostatic antibiotics, in *Vibrio cholerae*. We propose a molecular mechanism for the CBASS-antifolate interaction based on onset of cyclic-oligonucleotide production by the nucleotidyltransferase DncV upon folate depletion by antifolates. CBASS-antifolate interaction is specific to CBASS systems with closely related nucleotidyltransferases and similar folate binding. Altogether, our findings illustrate that toxic modules, such as the antiphage defense CBASS system, can dramatically impact antibiotic activity, and open the possibility that endogenous metabolites could also act as triggers/silencers of toxic modules under stress beyond antibiotic treatment, such as during phage infection, biofilm formation or disease environments.

## Main

Toxic modules, such as TA or abortive infection (Abi) systems, are involved in a wide range of cellular processes, e.g. stabilization of genetic elements, stress response, persister formation, or phage defense^4,7–9^. Due to their toxic nature, TA systems have been previously suggested as potential druggable targets, for instance by designing peptides that disrupt TA complexes, thereby unleashing toxicity^6,10,11^, among others^1–3,5,12^. Such strategies focus on targeting the TA itself to trigger its toxicity. However, antimicrobial activity is not only influenced by compound primary target, but also by proteins and pathways which are apparently not related to it^13,14^. In this study, we use a phenotypic screening approach, which is blind to whether a compound directly or indirectly targets the toxic module, to find compounds of which antimicrobial effect changes in the presence/absence of the toxic module. We selected the antiphage Abi CBASS system from the pandemic strain *V. cholerae* El Tor N16961 as a representative toxic module. *V. cholerae*’s CBASS is a well characterized example of the numerous newly discovered antiphage defense systems, containing several new types of potentially toxic modules with novel enzymatic functions^15–22^.

Abi systems are the last bacterial line of defense against bacteriophages, where an infected cell triggers its own death to limit phage replication^8^. They are typically (but not exclusively) encoded together with other antiphage defense systems in so-called defense islands, prophages, and mobile genetic elements, and their occurrence is strain-specific^15,20,23–25^. CBASS systems are related to the human antivirus cGAS-STING^17,26,27^, and are widespread in bacteria and archaea^18^. They contain at least two proteins: a sensor protein (generally termed nucleotidyltransferase, CD-NTases) that senses phage infection and produces a cyclic signaling molecule, and an effector protein that exerts the bacterial suicide^17,28^. In particular, the CBASS of *V. cholerae* El Tor N16961 is a four-gene system coding for a cyclase as sensor (DncV), a phospholipase as effector (CapV), and two accessory proteins (Cap 2 and Cap3), which prime DncV activity (Fig. 1a)^17,29,30^. DncV produces cyclic GMP–AMP (cGAMP), the signaling molecule that activates CapV, thereby causing lysis of phage-infected cells, and preventing further propagation of the infection^17,31^. CBASS from *V. cholerae* El Tor N16961, and in particular its DncV protein, is well characterized regarding its structure and cyclase activity^32,33^, and it has also been suggested to play a role in pathogenicity, biofilm formation, and motility of this bacterium^30^.

**Figure 1:**
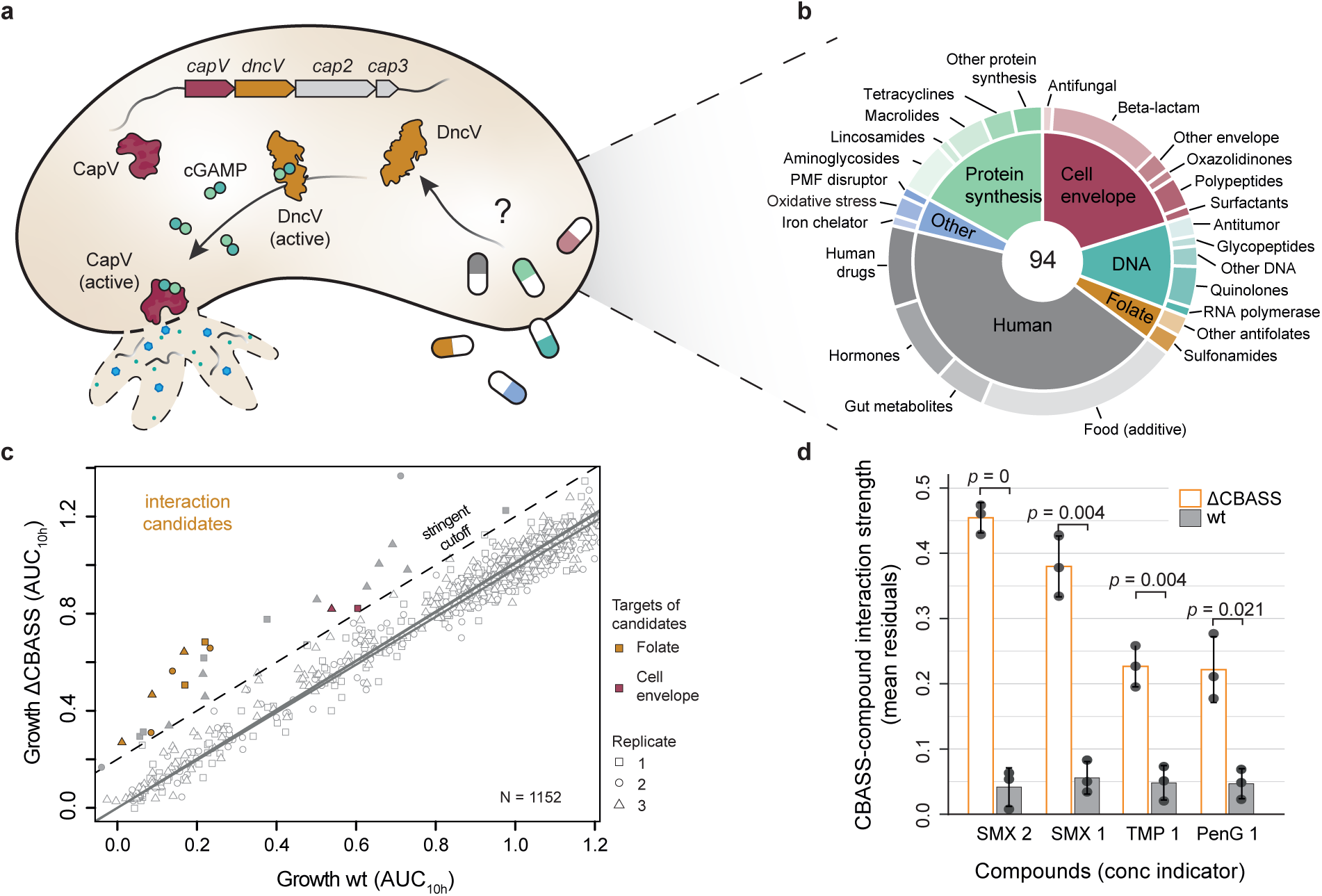
CBASS influences antimicrobial activity in *V. cholerae* el Tor N16961. **a)** Schematic representation of the components of the CBASS system of *V. cholerae* N16961. The CBASS operon encodes the phospholipase CapV (red) which is activated by the signalling molecule cGAMP (blue-green) produced by the cyclase DncV (yellow), thereby inducing cell death by lysis. **b)** Composition of the chemical library tested in this study, classified by target (in the case of antimicrobials, inner ring) and compound class (outer ring). Tested compounds cover all major antibiotic classes, plus “human targeted” compounds, such as endogenous hormones, food compounds and approved non-antibiotic drugs. Complete list and experimental details provided in Supplemental Table 1 and Extended Fig. 1. **c)** Comparison of treatment effects on the growth of wild-type V. cholerae and ΔCBASS strains. Experiments were done in triplicate, and each compound was tested in 4 concentrations. Growth is represented as area under the curve after 10h of treatment (AUC_10h_) for all tested conditions and replicates, with wild type on the x-axis and ΔCBASS on the y-axis. CBASS deletion did not affect treatment outcome in most conditions (open grey symbols). Data points that deviate from the line of best fit (residuals) by more than the a stringent cut-off of 0.2 – interaction candidates - are shown as solid symbols. The stringent cut-off (0.2) to assign CBASS-compound interactions was selected based on the variability observed for the three wild type replicates (Methods and Extended Data Fig. 1b). When ΔCBASS-wt residuals are significantly different from the wt-wt residuals, symbols are colour coded according to the target of the respective treatment from Fig 1b. **d)** CBASS-compound interaction strength as measured by mean ΔCBASS-wt residuals. Wt-wt residuals as shown for comparison. Only interaction candidates with Δ CBASS-wt residuals significantly different from wt-wt residuals are shown. SMX = sulfamethoxazole, TMP = trimethoprim, PenG = penicillin G.

### CBASS impacts resistance and synergy between antifolate antibiotics

We sought out to systematically investigate the impact of CBASS on antimicrobial activity by assessing bacterial growth of the wild-type *V. cholerae* and CBASS-operon deleted (ΔCBASS) strains in the presence of 94 small molecules, including antibiotics, human drugs, human endogenous metabolites and food additives (Fig. 1b, Supplementary table 1). Briefly, each strain was grown in the presence of the individual compounds at four different concentrations, and growth (OD_600 nm_) was frequently monitored for ∼12 h (ED Fig. 1a & b). The screen was performed in triplicates, with a Pearson correlation coefficient of 0.98 (ED Fig. 1c). In general the presence of CBASS did not change antimicrobial activity (Fig. 1c). This is in agreement with chemical genomic studies in bacteria, where bacterial fitness is highly robust to single gene deletions and compound stress^13^. Nonetheless, in few cases, CBASS modulated the activity of specific compounds, including the well-established antifolate antibiotics sulfamethoxazole (SMX) and trimethoprim (TMP), as well as of the *β*-lactam antibiotic penicillin G (Fig. 1d, ED Fig. 1d). In all cases, the ΔCBASS strain was more resistant to the antibiotics, suggesting that CBASS is relevant for their activity against this *V. cholerae* strain. With this initial observation, we establish that toxic modules can interact with established antibiotics, and change their antimicrobial effect.

We decided to further investigate the functional interaction between CBASS and antifolates (CBASS-antifolate interaction). Antifolates inhibit folate biosynthesis, thereby hindering the synthesis of DNA and RNA essential precursors^34^. They are used as antimicrobials, but also as antimalarials and anti-cancer drugs^34^. In bacteria, SMX and TMP inhibit enzymes of the folate pathway (dihydropteroate synthase and dihydrofolate reductase, respectively), and are most commonly used in combination (co-trimoxazole) to treat a wide range of infections^34–36^. We first validated our observation from the initial screen by re-evaluating antimicrobial activity of SMX and TMP against wild-type *V. cholerae* and ΔCBASS. As expected, we observed that the ΔCBASS strain grows better than the wild type at the same given concentration, particularly strong in the case of SMX (Fig. 2a). This prompted us to test whether *V. cholerae* ΔCBASS strain would be more resistant to both antifolates than wild type. Indeed, by determining the minimum inhibitory concentration (MIC), we observed that ΔCBASS is more resistant to SMX and TMP by a factor of ∼3- and ∼2-fold, respectively (Fig. 2b). Next, we assessed the impact of CBASS on the combinatorial treatment, because antifolates are often used together in a strong synergistic combination^35,36^. We used checkerboard experiments to obtain a response surface to the drug combination (isobologram). A strong synergy is maintained for both strains, wild type and ΔCBASS, with pronouncedly concave isoboles – lines of equal growth (Fig. 2c). Nonetheless, the synergy between the two antibiotics is weaker in the presence of CBASS. This is directly recognizable from the surface response (wild type isoboles less concave than ΔCBASS), and confirmed by the Loewe fractional inhibitory concentration index (FICi)^37,38^ (Fig. 2c, ED Fig. 2d, FICi<1 indicates synergy). Importantly, despite the synergy strength being enhanced in ΔCBASS, higher absolute concentration of both antibiotics (> 2-fold) is still required in combinatorial treatment to achieve a comparable inhibition to the wild type (Fig. 2d). Thus, so far we show that CBASS is not only modulating sensitivity to antifolates in *V. cholerae*, but also interfering with the strength of their synergistic interaction. Further investigation will be needed to fully understand how CBASS can weaken one of the most conserved antibiotic synergies. Here we limit the scope to investigate CBASS interaction with single antifolates, TMP or SMX.

**Figure 2:**
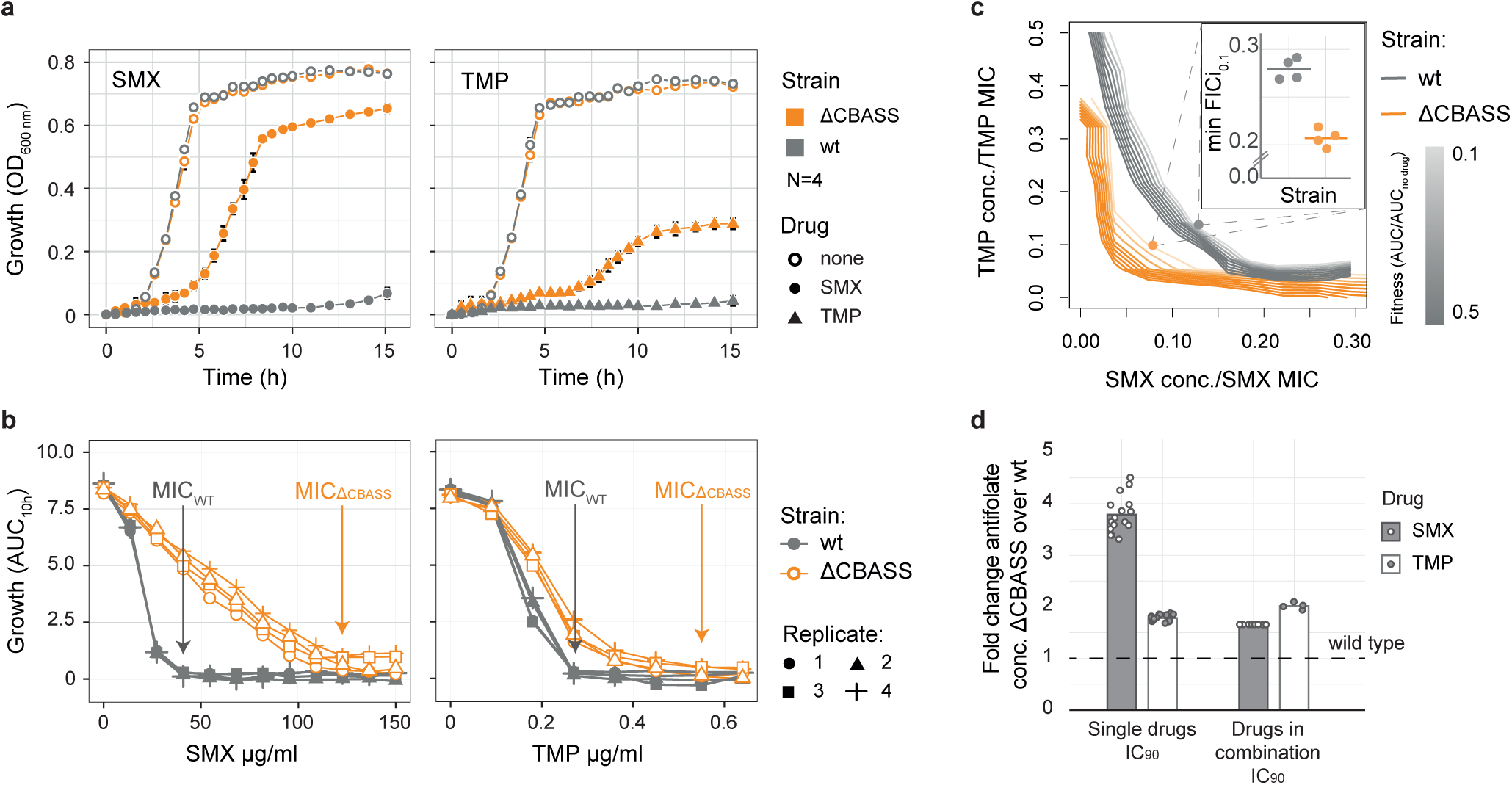
CBASS impacts resistance and synergy to antifolate antibiotics in *V. cholerae*. **a)**ΔCBASS grows better than wild-type *V. cholerae* in the presence of antifolates. Growth over time of wild-type V. cholerae (grey) and ΔCBASS (orange) cells treated with 41 µg/ml SMX (left panel), or 0.3 µg/ml TMP (right panel). **b)** ΔCBASS is more resistant to antifolates than wild-type V. cholerae. Minimum inhibition concentration (MIC) curves for SMX (left panel), or TMP (right panel). Growth of wild-type *V. cholerae* (grey) and ΔCBASS (orange) at various antibiotic concentrations was determined as AUC_10h_ and plotted against the respective antibiotic concentrations. Arrows point to MIC of wild type (grey) and ΔCBASS (orange). Results from 4 replicates are shown. Replicate correlation shown in ED Fig. 2a. **c)** CBASS impacts the synergy between antifolate antibiotics. Isobologram of SMX-TMP combination treatment shows stronger synergistic drug interaction in ΔCBASS (orange) compared to wild type (grey). Concentrations used in the checkerboards are shown relative to their respective single-drug MIC_95_. One exemplary replicate of four per strain is shown (replicate correlation shown in ED Fig. 2b), with isoboles representing concentration pairs of equal fitness. MIC curves were obtained for identical growth conditions (384 instead of 96 well-plates), shown in ED Fig. 2c. Panel insert shows the minimum fractional inhibitory index (min FICi_0.1_) along the 0.1 fitness isobole for all replicates (ED Fig. 2d). **d)** Δ CBASS requires higher antifolates concentration not only alone, but also in combination, when compared to wild-type *V. cholerae*. Comparison of SMX and TMP impact on growth as single or double treatment on ΔCBASS growth relative to wild-type *V. cholerae* (dashed-line). Concentration values to achieve 90% growth inhibition are represented as fold change relative to the wild type response to the respective drug.

To establish specificity of the CBASS-antifolate interaction regarding the toxic module, we tested whether a second antiphage system encoded in *V. cholerae* N16961 – Zorya – also influences antifolate activity. Zorya is a 4-gene system, also commonly found in defense islands. Similar to CBASS, it has been shown to confer phage protection, but its function and components are less well understood^15^. In analogy to our approach to uncover the functional CBASS-antifolate interaction, we generated a *V. cholerae* N16961 ΔZorya, and challenged it with SMX or TMP. As opposed to CBASS, antifolate activity is not changed between the wild type and ΔZorya (ED Fig. 2e & f), suggesting specificity of CBASS-antifolate interaction in *V. cholerae*. Even though this observation alone cannot completely exclude a Zorya-antifolate interaction, it confirms that the influence of antiphage defense systems on antimicrobial activity is most likely specific to compound – system – testing-conditions.

### Molecular insights of CBASS-antifolate interaction in vivo

To start gaining molecular insights into the CBASS-antifolate interaction, we asked which components of the four-gene CBASS system are required to enhance antifolate activity. We measured the MIC_90_ of SMX of each individual gene deletion mutant or, in case of DncV, a point mutation that renders the cyclase inactive (DncV_AIA_)^32^. Deletion of CapV or inactivation of DncV (DncV_AIA_) alone are sufficient to recapitulate the phenotype of the deletion of the entire operon, and make *V. cholerae* resistant to SMX (Fig. 3a). This shows that both, cGAMP production by DncV, and phospholipase activity of CapV are essential for the CBASS-antifolates interaction mechanism. While deletion of *cap3* did not affect sensitivity to SMX, deletion of *cap2* causes a mild increase in SMX MIC_90_, albeit not to the extent of either ΔCBASS, Δ*capV*, or DncV_AIA_ (Fig. 3a). Despite its reduced extent, this is a significant effect, indicating that Cap2 activity is important for CBASS function in the context of antifolate activity, in addition to its previously shown relevance for anti-page defense^29^.

**Figure 3:**
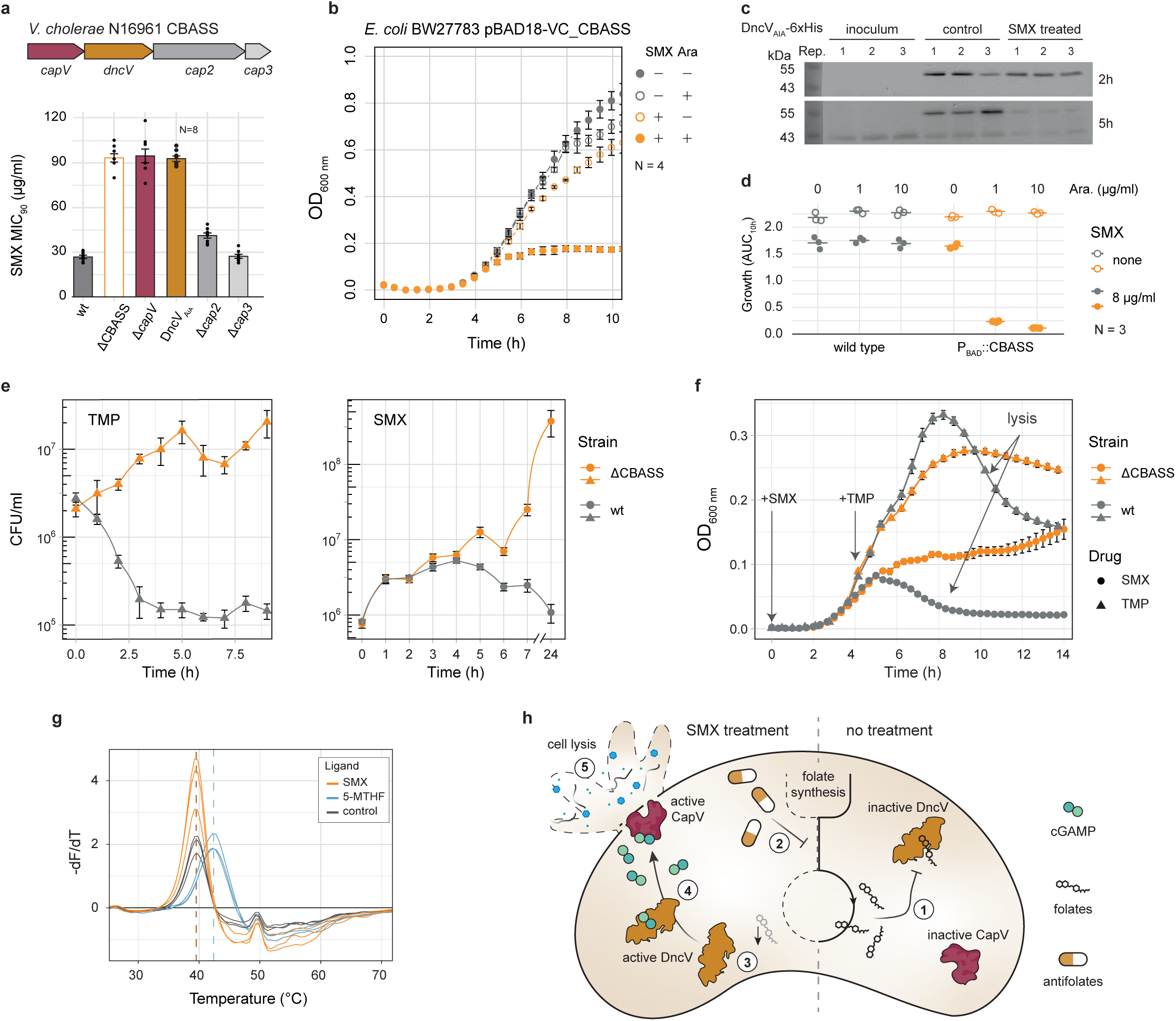
Molecular insights of CBASS mechanistic interaction with antifolate antibiotics. **a)** Intact DncV and CapV are essential for CBASS-antifolate interaction. SMX minimum inhibitory concentration (MIC_90_) for wild-type *V. cholerae* (wt) and mutants of individual CBASS genes. **b)** CBASS-antifolate interaction remains upon CBASS heterologous expression. Mean growth curves (OD_600 nm_) of *E. coli* BW27783 containing pBAD18-VC_CBASS, where CBASS expression is controlled by arabinose. SMX antimicrobial effect (orange symbols) is enhanced in the presence of arabinose (full versus empty orange symbols). SMX and arabinose concentrations used were 14 and 0.7 µg/ml, respectively. Experiments were done in quadruplicates. **c)** SMX treatment does not increase CBASS levels. Immunoblot analysis of DncVAIA-6xHis (∼50 kDa) using anti-His antibodies. Shown are three replicates of non-treated cells, and cells treated with 200 µg/ml SMX for 2h (upper panel) or 5h (lower panel) post SMX-addition. Inoculum in both panels is the same. Total protein loaded per lane was controlled by Coomassie stain (Supplementary Fig. 1). **d)** CBASS overexpression exacerbates the CBASS-antifolate interaction. Growth (AUC_10h_) of wild-type V. cholerae (wt, grey) and CBASS-inducible mutant where the endogenous promoter region is replaced by araBAD promoter of pBAD18 (orange). Cells were treated with 0, 1, or 10 ‰ (w/v) L-arabinose and either 0 (empty circles) or 8 (full circles) µg/ml SMX. Experiments were done in triplicates. **e)** CBASS enables killing by antifolates. Colony forming units (CFU) of wild-type *V. cholerae* (grey) and ΔCBASS (orange) treated with 200 µg/ml SMX (left panel) or 1 µg/ml TMP over at least 9h are shown. Experiments were done in biological triplicates. **f)** Wild-type *V. cholerae* cells containing CBASS lyse upon antifolate treatment. Mean growth curves (OD_600 nm_) of wild-type *V. cholerae* (grey) and ΔCBASS (orange) treated with 200 µg/ml SMX, or 2 and 6 µg/ml TMP for wild type and ΔCBASS, respectively. Experiments were done in biological triplicates. **g)** Thermal shift assay confirms binding of 5-MTHF to DncV, but no direct SMX-DncV interaction. Shown are derivate melting curves (-dF/dT) over the applied temperature gradient of purified DncV treated with a ∼80-fold molar excess of 5MTHF (blue) or SMX (orange), and a buffer control (grey). The maximum of the curves indicates the melting temperature of DncV in each treatment, highlighted by dashed lines in the respective colours. N=3. h) Proposed mechanistic model for the molecular mechanism of CBASS-antifolates interaction. 1 – in its normal state, DncV is inhibited by folate species bound to the allosteric pocket. 2 – Upon treatment with antifolates, the folate pool is depleted, leading to 3 – a shortage of folate molecules necessary to repress DncV. 4 – DncV becomes active, thereby producing cGAMP, which activates the phospholipase CapV. 5 – Enzymatic activity of CapV leads to cell death by lysis. Steps 1, 2, 4 and 5 were previously shown^17,30–32^. Step 3 is novel, and the key to the molecular mechanism of CBASS-antifolates interaction.

Our finding that deletion of CapV or inactivation of DncV completely abrogated the interaction with antifolates, indicates that activity of the system is crucial for the CBASS-antifolate interaction. This result could be further confirmed when we expressed CBASS in an heterologous host (*Escherichia coli* BW27783), thereby eliminating all factors brought in by genetic context, and again observed that CBASS renders *E. coli* highly sensitive to SMX (Fig. 3b). Hence, our observations support an activity-based mechanism, where CBASS is not only necessary, but also sufficient for the CBASS-antifolates interaction.

It has been previously observed that over-production of DncV leads to increased intracellular cGAMP levels *in vivo*, which in turn activates CapV-dependent cell lysis^31,39^. Thus, we hypothesized whether SMX induces higher DncV levels, thereby triggering CBASS toxicity and explaining higher SMX sensitivity of the wild type. We generated a scarless C-terminal 6xHis-tagged version of DncV expressed from the original chromosomal locus. To avoid DncV toxicity interfering with the measurement, we used the inactive DncV mutant DncV_AIA_. SMX treatment did not increase DncV levels, and, if anything, led to lower protein levels at later time points (Fig 3c). Interestingly, DncV was also barely detectable in untreated cells during late stationary phase (Fig. 3c; inoculum). This suggests that DncV protein levels might decrease during less-favorable growth conditions, and that its decreased levels in the presence of SMX are a consequence of limited growth caused by the antibiotic, rather than a direct effect of the antibiotic on CBASS. We reasoned that this effect could generate an inhibition feedback cycle, where decreased growth by SMX prevents an even stronger CBASS contribution to its antimicrobial activity. We chromosomally expressed CBASS in *V. cholerae* under an inducible promoter to disrupt its native regulation and break the potential feedback cycle. We observed an increased SMX sensitivity compared to the natively regulated system, already with moderate/low CBASS expression (Fig. 3d). Hence there is a directly link between CBASS and SMX activities, which, if anything, gets ameliorated by the native regulation of CBASS levels.

Since the effector of *V. cholerae* CBASS is a phospholipase, a direct consequence of CBASS activity is cell death by lysis^31^. While antifolates typically act bacteriostatically (only inhibit bacterial growth, but do not kill)^40^, we observed cell death of wild-type *V. cholerae* after treatment with SMX or TMP, particularly stronger in the case of TMP (Fig. 3e). Cell death is completely CBASS dependent, as the antibiotics have only mild bacteriostatic activity against ΔCBASS (Fig. 3e). Finally, loss of optical density of the culture happens simultaneously to cell death for both antibiotics, which is symptomatic for cell lysis (Fig. 3f). Importantly, the bacteriostatic effect on ΔCBASS remained when we used higher TMP concentrations to compensate for its higher resistance (Fig. 3f). Altogether, our results support a model where antifolates trigger CBASS activity, leading to cell death by lysis. To our knowledge, this is the first time that molecular evidence is provided as basis for killing activity of otherwise bacteriostatic antibiotics.

DncV activity has been shown to be allosterically inhibited by folate species *in vitro*, particularly by 5-methyl-tetrahydrofolate (5-MTHF)^32^. Indeed, we could also observe a thermal shift of purified DncV (from 39.5 to 42.4°C) upon incubation with 5-MTHF, corroborating a protein-ligand interaction (Fig. 3g). In addition, we failed to obtain a thermal shift with SMX, rendering it unlikely that SMX directly binds the toxic module component DncV (Fig. 3g). Thus, we hereby propose that antifolate treatment triggers CBASS activity *in vivo* by depleting the folate pool^41^, and thereby releasing DncV allosteric regulation. Consequent cGAMP production activates CapV, which causes cell lysis by hydrolysis of the cytoplasmatic membrane (Fig. 3h). This not only leads to an exacerbation of antifolates antimicrobial activity, but also enables killing by the antifolates. Interestingly, in the context of supporting that 5-MTHF is in fact inhibiting CBASS activity, it has been shown that antifolate treatment increases cGAMP production when DncV is heterologously overexpressed in *E. coli*^32^. However, the authors did not investigate further cellular consequences of their observation, which we now show to be extremely severe at CBASS endogenous levels in its native host: DncV enzymatic activation by antifolates impacts resistance, tolerance and even the extent of synergy between antifolates in *V. cholerae* El Tor N16961.

### CBASS-antifolates interaction is specific to CBASS systems with closely related CD-NTases

CBASS is one of the most prominent Abi systems. Almost 6,000 CBASS operons have been annotated across 15,000 bacterial and archaeal strains^42–44^. Even though the operon composition changes across systems, namely the nature of the effector and presence of accessory proteins like Cap2 and Cap3, they always contain a CD-NTase^28^. Since DncV is the key enzyme for interaction with antifolates, we sought out to explore how general is the CBASS-antifolate interaction across the wide genetic landscape of CD-NTases. We first confirmed that CBASS-antifolate interaction is conserved in the closely related *V. cholerae* strain A1552, which contains an identical CBASS system to *V. cholerae* N16961 (ED Fig. 3a). Next, we built a phylogenetic tree of all annotated CD-NTases, which recapitulated the ∼10 reported clusters (Fig. 4a) by previous studies^42,43^, and selected four other CBASS systems for querying their potential interaction with antifolates: one with similar CD-NTase to the one from *V. cholerae* N16961 – *E. coli* TW11681 (CBASS system 2), and three others phylogenetically more distant – *E. coli* 3234/A, *E. coli* TW11681 (CBASS system 1) and *E. coli* U5/41 (Fig. 4a & b). The phylogenetic distance of each protein to DncV is well recapitulated by difference in the 3D structure of the proteins, as predicted by AlphaFold^45^ (Fig. 4b & c). Next, we individually expressed all CBASS systems in *E. coli* BW27783 to avoid potential confounding factors bound to different genetic contexts. We first assessed toxicity of each system by tuning its arabinose-inducible expression. Four out of the five CBASS systems (including the one from *V. cholerae* N16961) showed toxicity in *E. coli* BW27783 when overexpressed (Fig. 4d). The most distant CBASS system – that of *E. coli* U5/41 – showed no toxicity, precluding us from assessing the CBASS-antifolate interaction. We then evaluated potential CBASS-antifolate interaction of the four remaining systems by treatment with SMX (Fig. 4e). Since CBASS toxicity is immediately very high upon induction (Fig. 4d), the initial assessment of CBASS-antifolate interaction was done without addition of inducer (arabinose), making use of the leaky expression known to the arabinose promoter in the absence of glucose^46^. Using the CBASS from *E. coli* U5/41 as control, we found that, in addition to the system from *V. cholerae* N16961, also the system from *E. coli* TW11681 (2), with the closely related CD-NTase, shows a strong decrease in SMX MIC (Fig. 4e). The CBASS systems with more distant CD-NTases (*E. coli* 3234/A and *E. coli* TW11681 (1)) did not show convincing difference in SMX MIC. To exclude the possibility that low CBASS expression could mask a potential CBASS-antifolate interaction, we tested the effect of SMX over a range of CBASS induction in a checkerboard-like fashion (ED Fig. 3d). We were able to confirm that induction of the system from *E. coli* 3234/A did not increase SMX toxicity, while that of *E. coli* TW11681 (1) seems to very mildly do so (ED Fig. 3d). Altogether, our findings suggest that strong CBASS-antifolates interaction is specific to CBASS systems with CD-NTases closely related to DncV, such as that of *E. coli* TW11681 (2). The predicted protein structure of the CD-NTase from *E. coli* TW11681 (2) is almost identical to DncV, including the allosteric pocket where folates were found to bind to inhibit cyclase activity^32^ (Fig. 4f). The strong interaction is lost for systems with distant CD-NTases, where a similar allosteric pocket seems to be absent, thereby supporting our proposed model that antifolates trigger CBASS activity *in vivo* by depletion of folates.

**Figure 4:**
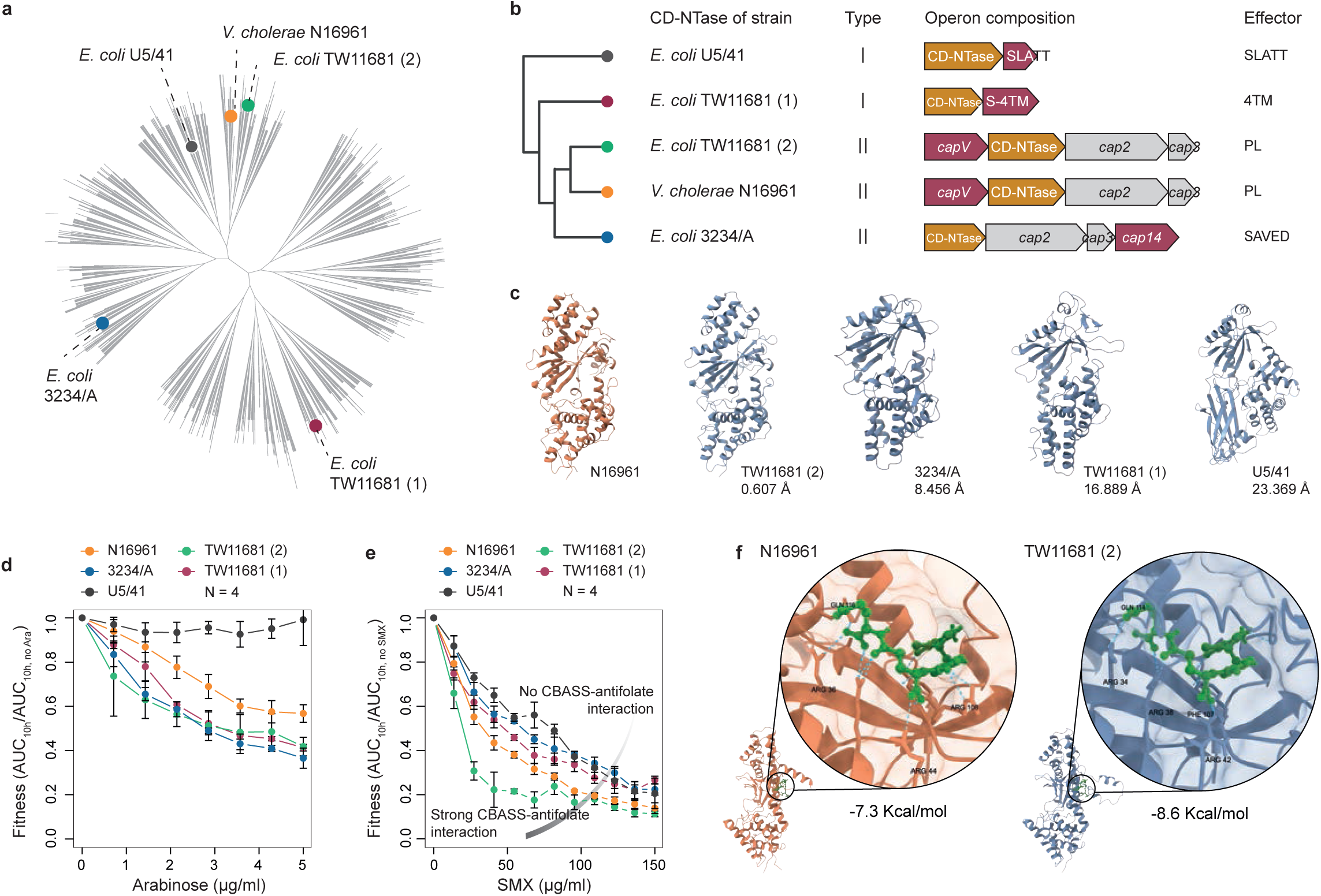
CBASS-antifolates interaction is specific to CBASS systems with closely related CD-NTases. **a)** Unrooted phylogenetic tree of 6056 previously predicted CD-NTases from CBASS systems_43_. Colored dots point to the systems selected for assessing conservation of CBASS-antifolates interaction across different CD-NTases. (Supplemental Table 3). **b)** Phylogram representing the genetic distance of the selected five CD-NTases, and schematic representation of the corresponding CBASS operon structure. CBASS system type, gene names and effector as per previous classifications^43,49,50^. SLATT = SMODS and LOG-Smf/DprA-Associating Two TM, S-4TM = SMODS-associating 4TM, PL = Phospholipase, SAVED = SMODS-associated and fused to various effector domains. **c)** 3D structures of the five CD-NTases. Crystal structure of *V. cholerae* DncV^32^ is shown in orange, and predicted structures for the remaining four CD-NTases (predicted with Alpha-Fold^45^) are shown in light-blue models. Parameters used for assessing quality of the structure predictions are shown in ED Fig. 3b & c. CD-NTase structures facing the predicted active site are shown. Similarity of predicted structures to DncV is given as average distance (Å) of all atom pairs of the Ca backbones as root mean square deviation (RMSD). **d)** Four out of the five tested CBASS systems become toxic against *E. coli* BW27783 when overexpressed. All CBASS systems were expressed using pBAD18 under arabinose induction. Growth (AUC10h) decreases with increasing concentration of arabinose for all systems, except for *E. coli* U5/41. Experiments were done in quadrupli-cates. **e)** Only the CBASS system containing the CD-NTase close to DncV shows strong CBASS-antifolate interaction. SMX minimum inhibitory concentration (MIC) curves are shown for *E. coli* BW27783 with leaky CBASS expression (no added arabinose) for each individual system. Because no toxicity was observed for the CBASS system of *E. coli* U5/41 alone (panel d), this strain is used here as a control as not-active CBASS, where CBASS-antifolate interaction is not expected. Experiments were done in quadruplicates. **f)** CD-NTase from *E. coli* TW11681 CBASS system 2 has an intact folate-binding allosteric pocket. Zoom in the allosteric pockets of CD-NTases from *V. cholerae* (left) and *E. coli* TW11681 (2) (right) bound to 5-MTHF. Free Gibbs energy calculated for docking 5-MTHF is shown below the corresponding protein.

In summary, our study provides evidence that bacterial toxic modules, such as the antiphage defense CBASS system, can strongly modulate the activity of well-established antibiotics. Furthermore, we shown CBASS does so as a cellular consequence of antibiotic activity – onset of CBASS activity upon depletion of folates by antifolate antibiotics, rather than directly interacting with the antibiotic – antifolates do not biochemically interact with CBASS. Thus, we speculate that other toxic modules could change antimicrobial activity through non-trivial mechanisms, a trait that is vastly unexplored. The presence of CBASS increases the sensitivity to antifolate antibiotics, interferes with their well-established and clinically relevant synergy, and enables killing by these antibiotics, which are otherwise bacteriostatic against *V. cholerae* ΔCBASS. Classical bacteriostatic antibiotics have recently been reported to become bactericidal in different strains and species – how this changes depending on the genetic background remains an open question^47^. Here we show evidence that toxic modules, such as CBASS, can provide a direct genetic basis for this phenomenon. We also propose that folate species act as molecular triggers for specific CBASS systems *in vivo*. So far, only phage-related molecular factors have been proposed to trigger Abi systems, like CBASS^48^. Thus, our findings suggest that small bacterial endogenous molecules could act as modulators of toxic modules *in vivo* upon stress, and potentially trigger their activity in different physiological contexts, such as phage infection, biofilm formation or disease environments. Finally, the specificity of CBASS-antifolate interaction to closely related CD-NTases opens the question of how are other CBASS systems regulated, and whether their activity is controlled by other endogenous metabolites in a similar fashion.

## Supporting information

Supplementary Figure 1: Coomassie gel used for control of total protein for Western Blot shown in Fig. 3.

## Acknowledgements

We thank Athanasios Typas (EMBL Heidelberg), Pedro Beltrao (ETH Zurich) and Chase Beisel (HIRI, JMU Würzburg) for providing feedback on the manuscript, and the Brochado lab members for discussions. We acknowledge K. Thormann for sharing the pNTPS138-R6K plasmid and *E. coli* WM3064, Gabriel Moncalian Montes for sharing *E. coli* BW27783, Ulrich Dobrindt for sharing *E. coli* 3234/a, and Kurt Hanevik for sharing genomic DNA from *E. coli* TW11681. We thank Lars Schönemann (JMU Würzburg) for support with DncV protein purification. This work was supported by JMU internal funding and the Emmy Noether program of the German Research Foundation to ARB (GO 3161/1-1), the GSLS Postdoc plus program funding at JMU to SB, and the Hector Research Career Development Award (HRCDA) 2020 by the Hector Fellow Academy to ARB. AJO and MA are supported by the HRCDA and GO 3161/1-1, respectively.

## Author contributions

SB and ARB conceived and designed the study. SB and MA performed the experiments. AJO did the CD-NTases phylogenetic analysis, structure and molecular docking predictions. ARB and SB analyzed the data and wrote the manuscript. ARB supervised the study.

---

**Extended Data Figure 1:**
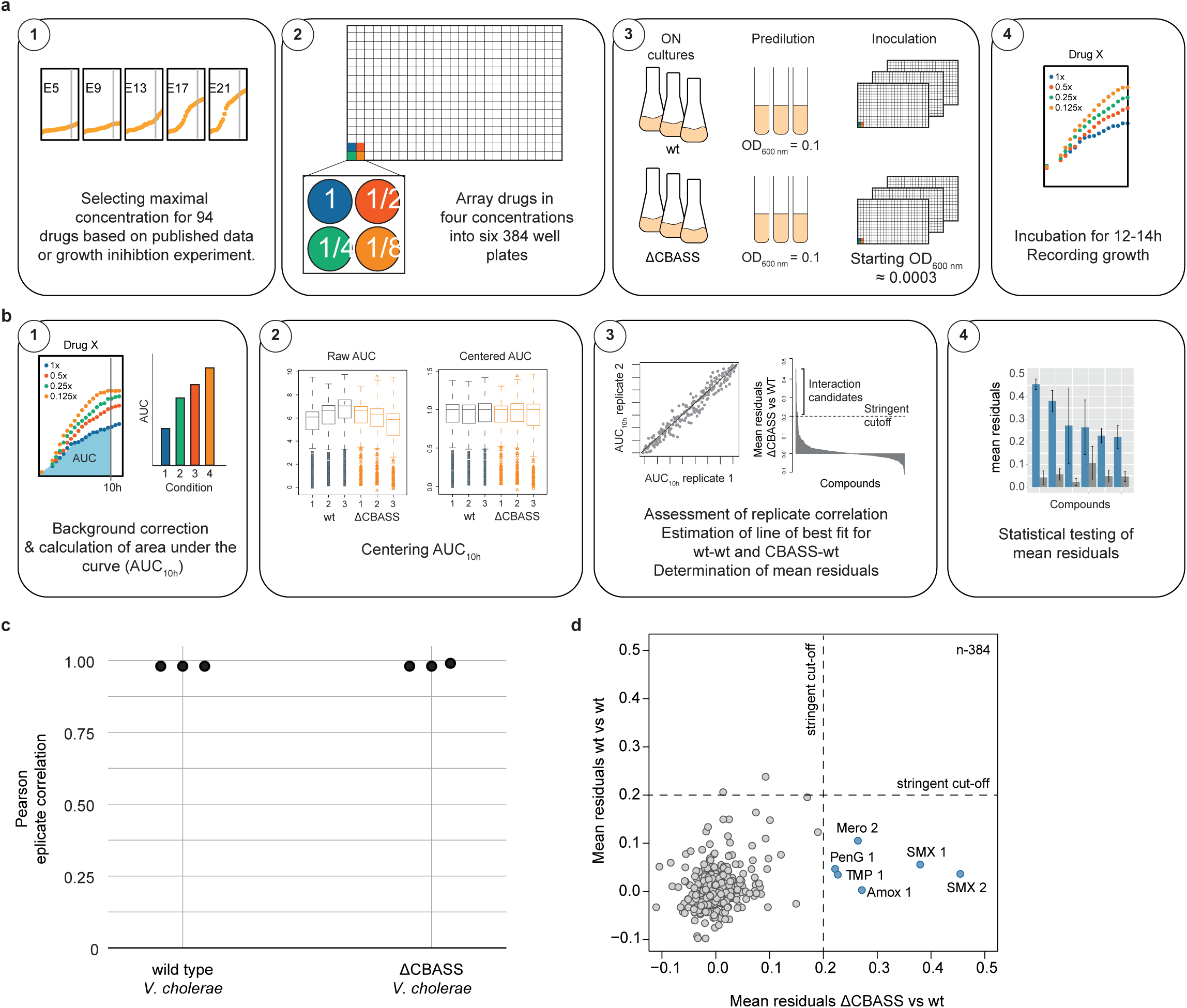
Schematic representation of the screen workflow and data processing. **a)** Schematic screen workflow. 1 – Maximal concentrations were selected based on literature and/or preliminary growth inhibition testing. In case of compounds without inhibition effect (e.g. by food additives), 500 uM was selected as maximal concentration. 2 – Compounds were dissolved in H2O, diluted to 0.5x, 0.25x, 0.125x concentration in LB, and arrayed into 384 well plates (40ul per well) using a Biomek i7 automated liquid handling workstation (Beckman Coulter). 3 – Inocula were prepared in triplicates from overnight cultures by pre-diluting to OD_600 nm_ = 0.5 before using a Singer rotor (Singer Instruments, UK) for inoculation - further ∼1:230 dilution step. 4 – OD_600 nm_ was recorded every 30 minutes for 12-14h. **b)** Schematic of data processing. 1 – OD_600 nm_ of LB (medium background) was subtracted from complete growth curves (OD_600 nm_ versus time). OD_600 nm_ of LB was estimated by the minimum OD_600 nm_ from each well. Area under the growth curve until 10h (AUC_10h_) is used as a as proxy for growth, and it was inferred by the trapezoid rule. 2 – AUC_10h_ of each well-plate (6 in total) was centered around 1 by division by the median of the respective plate. 3 – Replicate correlation was assessed by Pearson correlation. Compound-CBASS interaction scores were calculated to be the residuals of the line of best fit between the AUC10h of the ΔCBASS and the wild type for each replicate. 4 – t-test comparison between CBASS-vs-wt and WT-vs-WT with subsequent Benjamini-Hochberg correction for multiple testing to assign CBASS-compound interactions. **c)** Pearson replicate correlation between the triplicates of each strain. **d)** Highest 30 mean residuals from wild type-wild type analysis. Dark grey bars indicate the two treatments resulting in residuals >0.2. Black line highlights the cut-off value chosen for subsequent analysis. d) Scatterplot for comparison of mean residuals of wt vs. wt (y-axis) and CBASS vs. wt (x-axis). Each point represents the mean residuals of an given condition (compound, concentration) of the lines of best fit across three replicates. There are very few conditions for which wt vs. wt residuals (measure of noise) surpasses the stringent cut-off = 0.2. All conditions for which the CBASS vs. wt mean of residuals is higher than 0.2 are marked in blue. SMX = Sulfamethoxazole, TMP = Trimethoprim, Mero = Meropenem, PenG = Penicillin G, Amox = Amoxicillin.

**Extended Data Figure 2:**
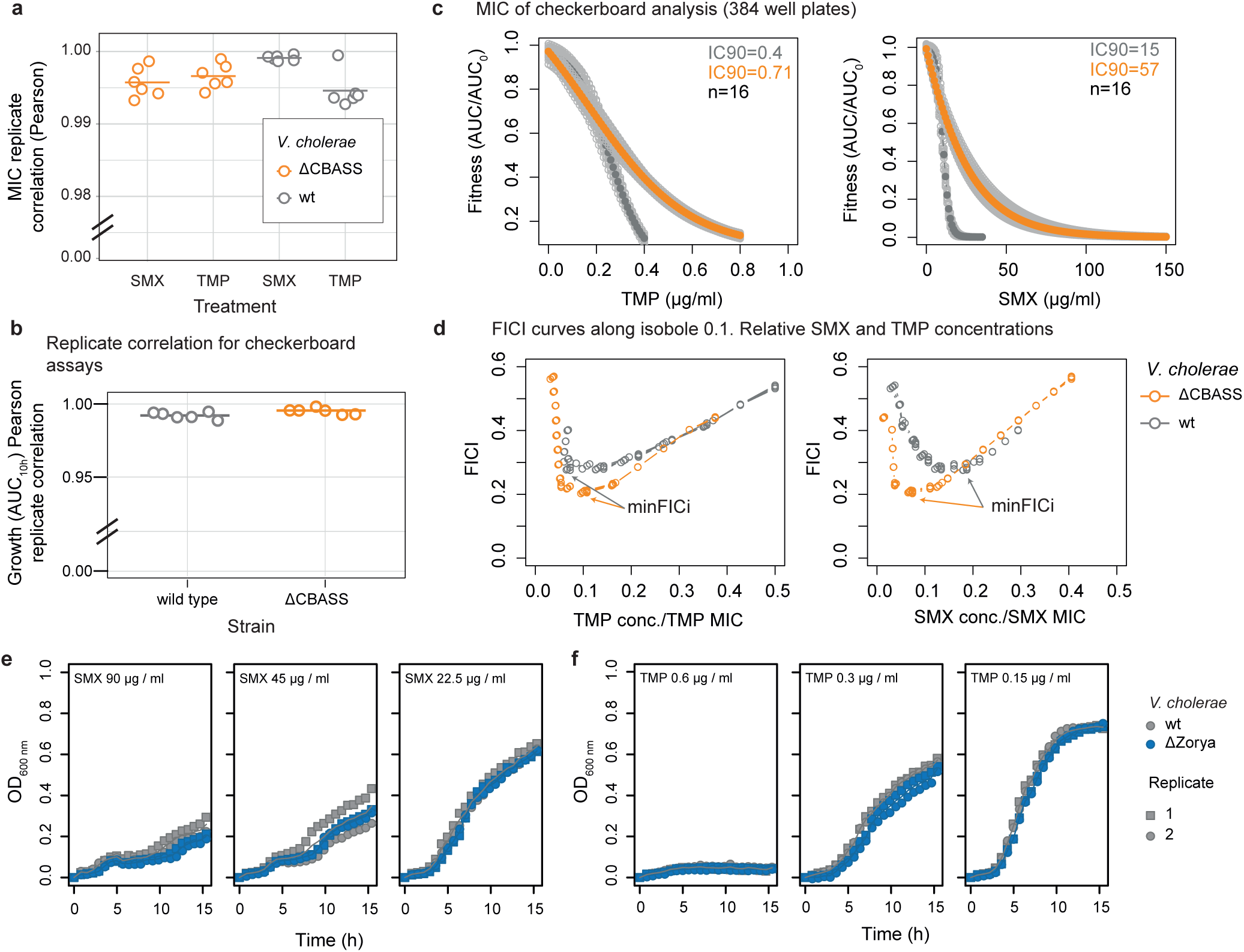
Impact of CBASS and Zorya in antifolates treatment of *V. cholerae*. **a)** Pearson replicate correlation of four MIC curves per strain and drug (data from Fig. 2b), obtained from growth experiments in 96 well plate. **b)** Pearson replicate correlation based on AUC_10h_ for checkerboard assays. **c)** MIC curves aquired in identical conditions as checkerboard experiment (384 well plates, N = 16) for TMP (left) and SMX (right). Gray and orange curves correspoind to wild-type *V. cholerae* and ΔCBASS, respectively. Drug concentrations are represented by the x-axis, while fitness is represented in y-axis. Fitness was calculated as the ratio between AUC_10h_ with and without added drug. **d)** Estimation of minFICi for isobole 0.1: FICis are calculated along drug concentrations normalized by their respective MICs, to determine the respective minimal minFICi_0.1_. **e & f)** Growth curves (OD_600 nm_) of wild-type V. cholerae (grey symbols) and *V. cholerae* deletion mutant of the Zorya operon (ΔZorya, blue symbols) at three e) SMX and f) TMP concentrations. Experiments were done in duplicates.

**Extended Data Figure 3:**
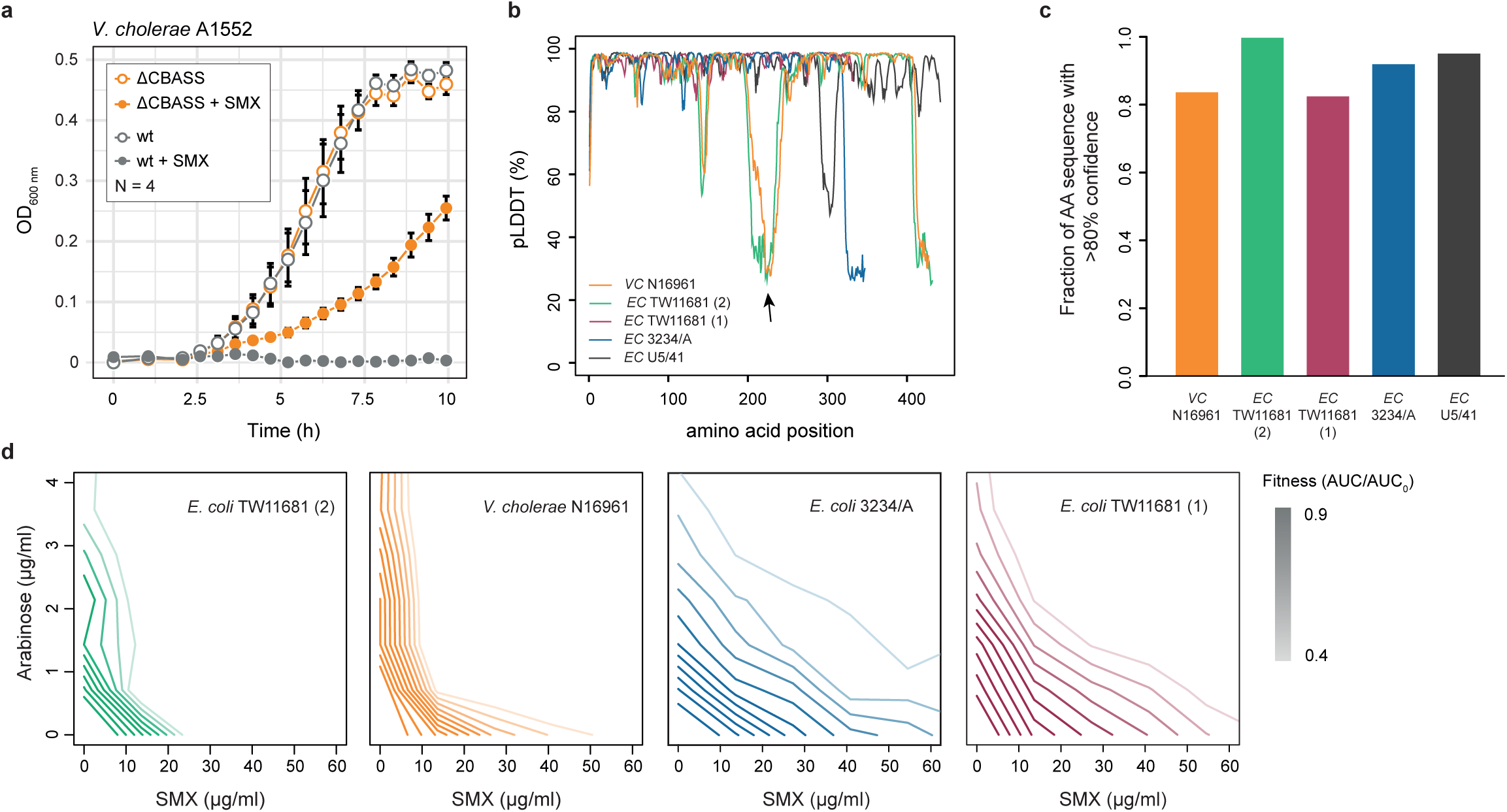
Assessment of CBASS-antifolate specificity across CD-Ntases. **a)** CBASS-antifolate interaction in *V. cholerae* A1552. Mean growth curves (OD_600 nm)_ of wild-type V. cholerae A1552 and its CBASS deletion mutant at 0 and 44 ug/ml SMX. Increased fitness upon CBASS deletion (open symbols) suggests that the CBASS-antifolate interaction is conserved between V. cholera N16961 and A1552. **b)** Predicted local distance difference test (pLDDT) of DncV and the four selected CD-NTases from additional CBASS systems as quality control parameter for structure prediction by AlphaFold46,52. DncV structure was also predicted as control. DncV structure was very accurately predicted in comparison with the crystal structure32 (RMSD = 0.655 Å). The lowest confidence was observed for unstructured C-termini and a region found to form a flexible loop in the crystal structure of *V. cholerae* DncV32 (AA 203–238, indicated by black arrow), and the similar CD-NTase from *E. coli* TW11681 CBASS system 2. **c)** More than 80% of all protein sequences have a high confidence score (>80%), indicating a high global performance of the structure prediction. **d)** Isobolograms of arabinose-SMX combination treatment confirm high CBASS-antifolate interaction (concave lines) for the *V. cholerae* and *E. coli* TW11681 (2) systems, and weak or no interaction for *E. coli* 3234/A and *E. coli* TW11681 (1). Shown is one exemplary out of four replicates per strain. Isoboles (lines) represent arabinose-SMX pairs with equal fitness. Fitness is calculated by dividing AUC10h of each arabinose-SMX pair by AUC_10h_ of the untreated samples.

## Methods

### Strains and growth medium

All strains used in this study are summarized in supplementary table 1. The wild-type epidemic *V. cholerae* El Tor N16961 was used throughout the study as a native CBASS-containing host (simply referred as wild type). *E. coli* BW27783 (K12 derivative) was used for heterologous expression of CBASS systems under titratable arabinose induction^1^. CBASS-containing strain *E. coli* U5/41 (DSM 30083) was purchased from DSMZ (Germany). Cells were cultured in Lysogeny Broth (LB Lennox) adjusted to pH 7.5 at 37°C (*E. coli*) or 32°C (*V. cholerae*). Growth medium was supplemented with kanamycin (50 µg / ml), carbenicillin (100 µg / ml), sucrose (10% (w/v)), or diaminopimelic acid (300µM), when selection or counter selection was required for strain construction. Compounds were purchased from Sigma-Aldrich-Merck, Switzerland.

### Strain construction

All plasmids generated for this study are listed in supplementary table 2. Scarless genomic manipulations were done with the suicide plasmid pNTPS-138-R6K for homologous recombination double crossover, using *sacB* gene counter-selection^2,3^. DNA fragments were generated using Q5 polymerase according to the manufacturer’s recommendation (New England Biolabs (NEB), USA). PCR oligo list is provided in supplementary table 3. All plasmids for genomic manipulation were assembled with Gibson assembly^4^, using EcoRV (NEB) for linearization of the backbone plasmid. Plasmid-based heterologous expression of CBASS systems in *E*.*coli* BW27783 was done using the pBAD18^5^.

### Compound screen

The compound screen was carried out by growing *V. cholerae* wild-type and ΔCBASS in 384 transparent well-plates (CatNo. 781162, Greiner-Bio One, Germany), in 40 µl LB Lennox medium. Each well contained a single compound. Water (no compound) was used as solvent control in 8 wells. All 94 compounds were purchased from Biomol (Germany), MP Biomedicals (Germany), or Sigma-Aldrich-Merck. Purchase details, solvents used for each compound, and concentrations used in the screen are listed in supplementary table 1. Maximum concentrations were selected to be close to MIC for compounds with antimicrobial activity. Otherwise, 500 µM was used as maximum concentration. Three 1:2 serial dilutions were done suing a Biomek i7 liquid handler (Beckman Coulter, USA), in order to obtain the four working concentrations. For workflow see Extended Data Fig. 1a. Cells were inoculated to OD_600 nm_ of ∼0.0003 from overnight cultures using a Singer Rotor (Singer Instruments, UK). Plates were sealed with transparent breathable membranes (Breathe-Easy®, Sigma-Aldrich-Merck) and incubated at 32 °C in a Cytomat 2 incubator (Thermo Scientific) with continuous shaking at 800 rpm. OD_600 nm_ was measured at regular 40 minutes intervals for up to 14 h in a Synergy H1 plate reader (Agilent, USA). Experiments were conducted in triplicates, so that 1152 complete growth curves were collected for each one of the two strains.

Data was analysed using R-language (version 4.2.2) as follows (schematic representation in Extended Data Fig. 1b): Growth curves (OD_600 nm_ versus time) were corrected by subtracting the OD_600 nm_ of LB. Area under the growth curve until 10h (AUC_10h_) was calculated to serve as a proxy for growth. This parameter was selected because it is robust to reflect growth defects originating from lag-phase, exponential growth or stationary phase. Data after 10h was not taken into account, as occasionally non reproducible late growth was observed, suggestive of selection of resistant mutants. These two initial steps of this analysis pipeline were re-used throughout this study to quantify bacterial growth. Next, AUC_10h_ of each well-plate was centered around 1 by division by the median of the respective replicate. Pearson replicate correlation of AUC_10h_ between all replicate pairs was calculated to assess data quality (Extended Data Fig. 1c). Compound-CBASS interaction scores were calculated to be the residuals of the line of best fit between the AUC_10h_ of the *β*CBASS and the wild type for each replicate (Fig. 1c). A stringent cut-off on compound-CBASS interaction scores (0.2) was assigned based on the direct comparison between the mean residuals of the line of best fit of *Δ*CBASS-*vs*-wt and wt-*vs*-wt (Extended data Fig. 1d). Function interaction with CBASS was accepted for compounds with the mean of their CBASS residuals was above the stringent cut-off (0.2), and the p-value of a t-test comparison with the WT-*vs*-WT residuals, after Benjamini-Hochberg correction for multiple testing, was less than 0.05.

### Quantification of antimicrobial activity of antifolates

Antimicrobial activity SMX and TMP against the different mutants was quantified at several levels: Minimum Inhibitory Concentration (MIC) was used to assess antibiotic resistance; checkerboard assays were used to quantify synergy between antifolates, and CBASS-antifolate interaction upon CBASS heterologous expression under arabinose inducing conditions; effect of single antibiotic concentrations on bacterial growth was used to measure differential antifolate effect across mutants; and Colony Forming Units (CFU) measurements were used to assess antibiotic killing. All experiments were done at least 3 times. Inoculum effect (drug potency increases with decreasing size of inoculum) of SMX and TMP has been described for several bacterial species^6–8^. For this reason, culture conditions, inoculum size and preparation were kept constant for direct comparison of antimicrobial activity between mutants, eg. MIC of wild-type vs *Δ*CBASS.

*MIC* measurements were done in 96-well plates using very similar conditions to those described for the compound screen. 11 (instead of 4) equally spaced serial concentrations of SMX and TMP were used to allow the generation of highly resolved drug dosage response curves (effect *vs* antibiotic concentration). SMX and TMP initial concentrations were selected according to the mutant at hand. Data analysis of the growth curves followed the initial steps as for the data analysis of the compound screen for calculating AUC_10h_. MIC calculation was done by fitting a logistic-like model to the drug dosage response curves using R-package *drc*.

*Checkerboard* assays were performed in 384-well plates, and bacterial growth was measured in 11 (horizontal dilution series), or 7 (vertical dilution series) equally-spaced concentrations of SMX and TMP, alone and in combination. Starting concentrations of SMX were 5 and 25 µg/ml for wild type and *β*CBASS, respectively. Starting concentrations of TMP were 0.2 and 0.3 µg/ml for wild type and *β*CBASS, respectively. Culture conditions, data acquisition and quantification of growth (AUC_10h_) was identical to the compound screen. Fitness was obtained by normalising AUC_10h_ of each well by the no-drug containing well. Synergy quantification was done based on highly resolved drug-interaction-surfaces (AUC_10h_ vs SMX vs TMT) following Loewe additivity model^9^ Lines of equal fitness (isoboles) were estimated by the contours derived from drug-interaction-surfaces. Fractional Inhibitory Concentration Index (FICi)^9,10^ was calculated along the isobole_0.1_ as follows:

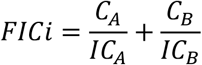

where C_A_ and C_B_ represent the concentration of the antibiotics A and B in combination for a given growth inhibition, IC_A_ and IC_B_ represent the concentration of the antibiotic A and B alone for the same growth inhibition. The minimum of the curves FICi vs SMX or TMP concentration – minFICi - was used as parameter for synergy quantification (Extended Fig. 2d).

*Single antibiotic concentrations* within dynamic range of MIC curves for each mutant were used to assess differential mutant behaviour. The experiments were carried out in 96 well-plates, using the same conditions as the ones used for MIC measurements. Quantification of growth (AUC_10h_) was identical to the compound screen.

*CFUs* were measured periodically in 15 ml cultures inoculated to an OD_600 nm_ = 0.005 for wild type and *β*CBASS for at least 10h, either with or without antifolates (biological triplicates). SMX (200 µg/ml) and TMP (1 µg/l) were added in the beginning to the experiment to avoid interference with inoculum effect. Each hour, 1 ml sample of each culture were retrieved, washed once to remove the antibiotic, and diluted in 1:10 steps. 10 µl of each dilution was plated onto LB plates, incubated at room temperature over night, and all dilution steps yielding individual colonies counted manually. The experiment with TMP was repeated for *Δ*CBASS with 3 µg/ml to exclude the possibility of potential concentration dependent killing activity.

*Lysis assay* was done in 96-well plates. Wild type and *β*CBASS were incubated in LB with or without SMX (200 µg/ml) or TMP (2 µg/ml and 6 µg/ml, respectively) for 14 h. Initial OD_600 nm_ was set to 0.005. OD_600 nm_ was measured every 40 minutes. SMX was added simultaneously with inoculum, to avoid an interference of the inoculum effect with assessment of lysis. TMP was added 4 hours after inoculation, as its premature supplementation did not allow sufficient growth to overcome the detection limit at OD_600 nm_ by the plate reader.

### Quantification of DncV_AIA_ levels by Western Blot

For immunoblotting of DncV_AIA_-6xHis, three cultures of *V. cholerae* N16961 *dncV*_*AIA*_-6xhis were grown overnight in LB at 32 °C. 2.5 ml of each overnight culture was pelleted by centrifugation (10 min, 4000 xg. 4 °C), the pellet flash-frozen in liquid nitrogen, and stored at -80 °C till further use. Per overnight culture, 2x 300 ml cultures were inoculated to OD_600 nm_ = 0.005 and incubated for 15 min at 32 °C before supplementing three replicates with SMX to a final concentration of 200 µg/ml. Untreated cultures were supplemented the equivalent amount of solvent. At 2 and 5 h post SMX supplementation, the OD_600 nm_ of SMX-treated and untreated cultures was measured. A volume containing an OD_600 nm_ > 3.5/ml was pelleted by centrifugation (see above), washed once in PBS, the supernatant removed, and flash-frozen in liquid nitrogen. All pellets were dissolved in PBS to a nominal OD_600 nm_ = 20, vortexed, the chromosomal DNA sheared by repeated passage through a 26 gauge needle, and boiled for 10 min. Total protein concentration was determined using the Pierce™ BCA Protein Assay Kit (Fisher Scientific, Germany) according to manufacturer’s instructions. 14µg protein per sample was adjusted to 1x Laemmli buffer supplemented with 2-Mercaptoethanol, boiled at 95 °C for 5 min, separated on a 10% SDS-PAGE and transferred to a PVDF membrane using the Trans-Blot® Turbo™ system (BioRad). His-tagged proteins were visualised using a primary anti-6xhis antibody (ArtNr: 11533923, Fisher Scientific), a HRP-coupled secondary antibody (ArtNr: 11515913, Fisher Scientific), and Pierce™ ECL Western Blotting-Substrate (Thermo Scientific) (Supplemental Figure). An SDS-PAGE with aliquots of the same samples was stained with Coomassie as a loading control.

### Protein purification and thermal shift assay of DncV

Purification of soluble C-terminal His-tagged DncV of *Vibrio cholerae* N16961 was done as previously described^11^. Thermal shift assays were performed using the GloMelt™ Thermal Shift Protein Stability Kit (biotium) and the Qiagen Rotor-Gene® Q PCR instrument according to the manufacturer’s recommendations. Each reaction contained a final concentration of 8.3 µM DncV-6xHis in thermal shift working buffer (50 mM Tris-HCl, pH8.0, 10 mM MgCl_2_, 100 mM NaCl), supplemented with 0.5x GloMelt™ Dye and either 0.653 mM Sulfamethoxazol or 5-Methyltetrahydrofolic acid (5MTHF), respectively, to yield a 1:78 protein: additive ratio, respectively. Melting curves were recorded in triplicates with ramps from 25 - 99°C and the melting temperatures calculated using the Rotor-Gene Q Software.

### CD-NTases multiple sequence alignment and phylogenetic tree construction

Protein sequences of known CD-NTases in bacterial and archaeal genomes (N=6056) as reported by Sorek *et al*.^12^ were downloaded from the JGI/IMG^13^ (https://img.jgi.doe.gov/) database in October, 2022. Sequences were aligned using MAFFT^14^ with a gap penalty of 2.0, offset value of 0.123 based on the BLOSUM62 matrix. Phylogenetic relationship was inferred with Fasttree^15^ using default parameters (neighbor-joining, no outgroups). Phylogenetic tree was visualized as an unrooted, tree with *ape* package in R^16^.

### CD-NTases protein structure prediction & similarity

Structures for CD-NTases of *E. coli* 3234/A, *E. coli* U5*/*41, *E*.*coli* TW11681(1), *E. coli* TW11681(2) were predicted by Colabfold^17^ (a wrapper for Alphafold2^18^). Since the crystal structure of DncV^11^ has been solved, we also predicted its structure as a form of positive control. Briefly, the amber force field was implemented to enforce accurate peptide geometry, while MMSeqs2 was used for multiple sequence alignment based on sequences from UniRef^19^. Predictions with the best local performance based on the average predicted local distance difference test (pLDDT), and sequence coverage score (global performance) were selected^18^. As a confidence measure, the pLDDT score of the four enzymes are comparable to the predicted DncV structure (Extended Figure 3b). In addition, the predicted DncV structure aligns well with its crystal structure (RMSD 0.665) hence all predicted structures are presumed to be of high quality.

The five non-DncV CD-NTases were superimposed on the DncV crystal structure (template, PDB: 4U0M^11^) using Pymol’s align function^20^. Root Mean Square Deviation, which is the average distance of all atom pairs of the Ca backbone between the query and template structures, was used as a metric for similarity. An increasing RMSD indicates an increasing sequence dissimilarity.

### Molecular docking

DncV and *E. coli* TW11681 (2) CD-NTase protein structures were prepared through energy minimization, charge neutralization and bond order reassignment on Autodock Tools 4^21^. For DncV, docking grids for the active (ATP) and allosteric sites were built using the coordinates of the co-crystallized ligands. To determine the coordinates of the docking grid center, we mapped the active sites using the primary sequences with MUSCLE^22^ and 3D superimposition of structures using Pymol^20^. All docking grids were restricted to a 40×40×40 Å^3^ size. Molecular docking of 5-MTHF was performed using Autodock 4.0 which implements a Lamarckian Genetic Algorithm to search for the most efficient conformation. After 10 resampling rounds, the binding predictions with the least free energy is selected for visualization and analysis^21^.

