## Supplementary figures and images for "The antiphage defense system CBASS controls resistance and enables killing by antifolate antibiotics in *Vibrio cholerae*"

### Supplementary Figure 1: Coomassie gel used for control of total protein for Western Blot shown in Fig. 3.

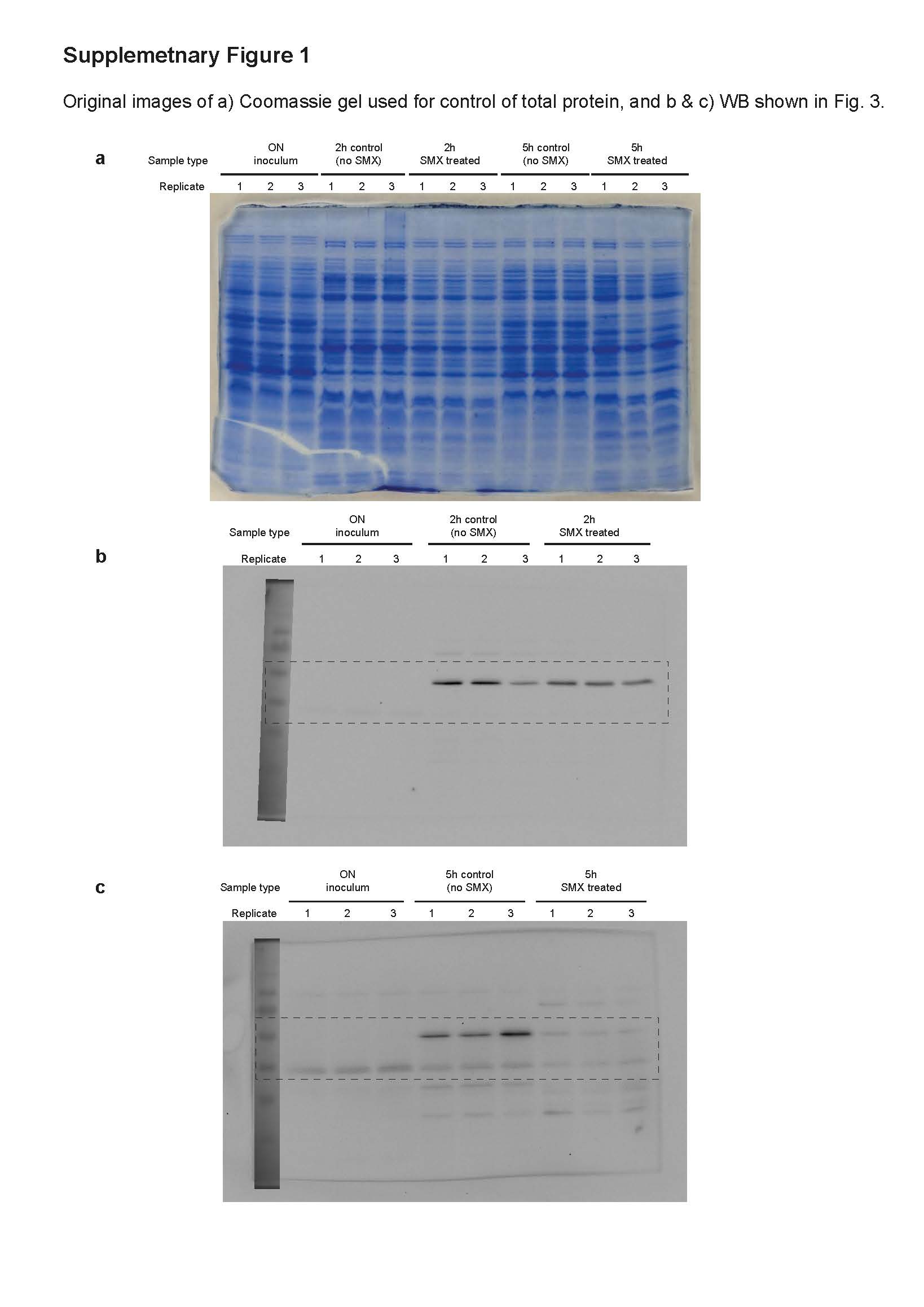
